# The stromal side of cytochrome *b*_6_*f* complex regulates state transitions

**DOI:** 10.1101/2023.08.08.552398

**Authors:** Alexis Riché, Louis Dumas, Soazig Malesinski, Guillaume Bossan, Céline Madigou, Francesca Zito, Jean Alric

**Author notes:** Corresponding author: Jean Alric.

## Abstract

In oxygenic photosynthesis, state transitions distribute light energy between Photosystem I and Photosystem II. This regulation involves the reduction of the plastoquinone pool, activation of the STT7 protein kinase by cytochrome *b*_6_*f* complex, phosphorylation and migration of Light Harvesting Complexes II (LHCII). Here we show the C-term of cyt *b*_6_ subunit acts on phosphorylation of STT7 and state transitions. We used site-directed mutagenesis of the chloroplast *petB* gene to truncate (remove L215*^b^*^6^) or elongate (add G216*^b^*^6^) the cyt *b*_6_ subunit. Modified complexes are devoid of heme *c*_i_ and degraded by FTSH protease, revealing that salt bridge formation between cyt *b*_6_ (PetB) and subunit IV (PetD) is key to the assembly of the complex. In double mutants where FTSH is inactivated, modified cyt *b*_6_*f* are accumulated but the phosphorylation cascade is blocked. We also replaced the arginine interacting with heme *c_i_* propionate (*R207K^b^*^6^). In this modified complex, heme *c*_i_ is present but the kinetics of phosphorylation are slower. We show that highly phosphorylated forms of STT7 are accumulated transiently after reduction of the PQ pool, and represent the active forms of the protein kinase. 96% protein coverage using phosphoproteomics showed 4 new phosphorylated peptides in the kinase domain of STT7. Phosphorylation of the LHCII targets is favored at the expense of the protein kinase, and the migration of LHCII towards PSI is the limiting step for state transitions.

**Significance statement:** State transitions are regulatory mechanisms that optimize the quantum yield of photosynthesis. *Chlamydomonas reinhardtii* is the choice organism to study this regulation at the molecular level. Our study describes an unprecedented mechanism of stereochemical changes at the Q_i_ site of the cytochrome *b*_6_*f* complex that trigger STT7 protein kinase activation through autophosphorylation.

## Introduction

In oxygenic photosynthesis, two photosystems PSII and PSI, connected by the cytochrome *b*_6_*f* complex, work in series to transfer electrons from a primary electron donor, water, to a terminal electron acceptor, NADP^+^. Electron transfer is coupled to proton translocation across the thylakoid membrane for generating a proton-motive force used for ATP synthesis. The reduction of CO_2_ into C3 compounds requires ATP and NADPH. State transitions optimize the yield of photosynthesis by distributing the mobile light harvesting complexes II (LHCII) between PSII and PSI. When the plastoquinone (PQ) pool is oxidized, dephosphorylated LHCII attach to PSII (State 1), and conversely when the excitation pressure on PSII is too high, giving an over-reduced PQ pool, LHCII are phosphorylated and migrate towards PSI (State 2). In *Chlamydomonas reinhardtii* the STT7 protein kinase contains a transmembrane helix and a stromal kinase domain that phosphorylates LHCII ^1^. The PBCP and PPH1 phosphatases, localized to the stroma of chloroplasts, dephosphorylate LHCII ^2^. Whereas PBCP and PPH1 are constitutively active, STT7 activity is regulated by a number of factors.

In the absence of cytochrome *b_6_f* complex ^3^, or when its lumenal PQ binding site Q_o_ is obstructed ^4–6^, state transitions are blocked. The transmembrane domain of STT7 interacts with cytochrome *f* and the Rieske subunit of cyt *b_6_f* ^7, 8^. The kinase domain of STT7 is on the other side of the thylakoid membrane where the stromal loop of cyt *b*_6_*f* subunit IV, between helixes F and G, allows for STT7 autophosphorylation *in vitro* ^9^.

In mitochondrial and bacterial cytochrome *bc*_1_, the cytochrome *b* subunit contains 8 transmembrane helixes and it is encoded by a single *petB* gene, see Figure 1A. In cyanobacteria and chloroplasts, cytochrome *b*_6_*f* complex binds a chlorophyll a and a β-carotene molecule, and the cytochrome *b* is split in cyt *b*_6_ and subunit IV (suIV), see Figure 1B, encoded by *petB* and *petD* respectively. While the split of cytochrome *b* into *b*_6_ and suIV leaves the coordination of the *b*-hemes and the quinone binding pocket Q_o_ unchanged ^10^, several structural rearrangements take place: helix H is lost and heme *c*_i_ occupies the Q_i_ site, covalently bound by C35*^b^*^6^.

**Figure 1.**
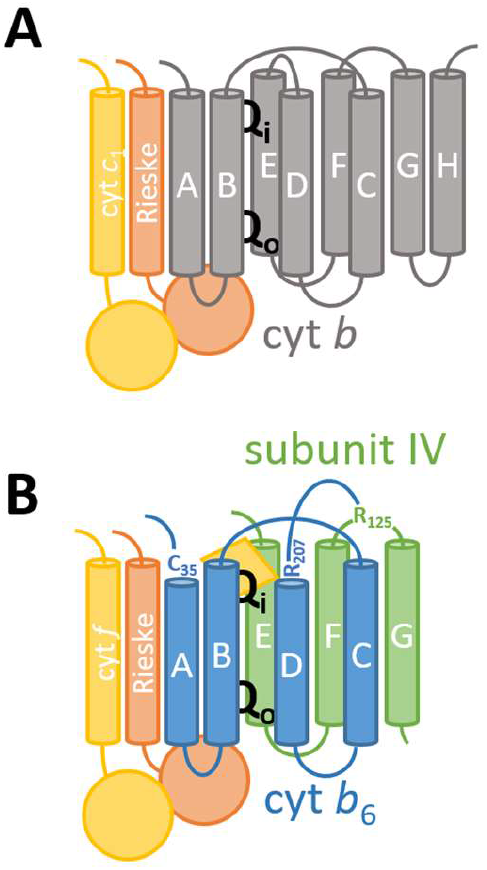
Comparison between (A) the mitochondrial cytochrome *bc*_1_ complex monomer and (B) the chloroplast cyt *b*_6_*f* complex monomer. In the latter, the cyt *b* is split into cyt *b*_6_ and subunit IV, helix H is missing, C35*^b^*^6^ binds heme *c*_i_ (yellow diamond) and helix D is not connected to E but to the inter helix F and G loop R125^suIV^. Cyt *b*_6_*f* also contains a β-carotene and a chlorophyll a molecule (not drawn).

In a previous work ^9^ we noted that the C-terminal end of the cytochrome *b*_6_ subunit forms a salt bridge with Arg125 of suIV. We showed R125^suIV^ is involved in a direct interaction with the STT7 protein kinase. Truncation or elongation of cyt *b*_6_ was done by removing the last Leu215 (*xL215^b^*^6^) or adding an extra Gly216 (*G216^b^*^6^). Amino acid R207*^b^*^6^, that interacts with the propionate group of heme *c*_i_ in cytochrome *b_6_f* crystal structure ^10^, was substituted with a similarly positively charged Lys in strain *R207K ^b^*^6^.

In this work we confirmed that the stromal region of cyt *b*_6_*f* is intimately connected to the STT7 protein kinase, not only at the level of suIV, but also through an interaction with the C-terminal part of cyt *b*_6_. In a variety of strains, we studied the kinetics of protein phosphorylation using PhosTag PAGE and western blot to propose a mechanistic model for state transition.

## Results

### Truncated or elongated versions of cyt *b*_6_ are degraded by FTSH

*C. reinhardtii* mutants were obtained by site-directed mutagenesis and chloroplast transformation of a *ΔpetB* strain with pWBA plasmids carrying an *aadA* resistance cassette and various mutated versions of the *petB* gene (for the complete list of strains used in this study see Supplementary Table S1). Transformants were selected on TAP medium supplemented with spectinomycin and sequenced to verify the presence of the mutated *petB* genes. While we successfully complemented two different *ΔpetB* strains with the pWBA plasmid carrying the WT *petB* gene, the mutants (*R207K*, *xL215* and *G216*) did not accumulate the cyt *b*_6_ subunit (Supplementary Figure S1) neither did they grow under photoautotrophic conditions. Therefore, cyt *b*_6_ C-terminal modifications specifically impede cyt *b*_6_*f* assembly and function. The mutant with an inactivated ATP-dependent zinc metalloprotease FTSH ^11^ (R420C substitution in strain *ftsh1-1*, kindly provided by Catherine de Vitry) was then used as a host strain for transformation with the pWBA plasmids carrying mutated or non-mutated *petB*. Transformants were selected on TAP medium containing spectinomycin. Genotyping by PCR showed homoplasmy was reached after > 3 months of restreaking under antibiotic resistance pressure, see Supplementary Figure S2.

Genetically modified strains were tested on plates and compared to reference strains (Figure 2). WT and *stt7-1* ^1^ grew on tris-acetate-phosphate (TAP) and minimal (MIN) media while the cyt *b_6_f*-less strain *ΔpetB* ^12^ required acetate. The control strain (*ftsh1-1* transformed with non-mutated pWBA plasmid*) ftsh*, and site-directed mutants of the *petB* gene *R207K^b^*^6^, *xL215^b^*^6^ and *G216^b^*^6^ were able to grow slowly under phototrophic conditions. Mutants *xL215^b^*^6^ and *G216^b^*^6^ have low photosystem II quantum yield ϕ_PSII_ and high F_M_’/F_M_ (see Materials and Methods for chlorophyll fluorescence), similarly to *ΔpetB*, while *ftsh* and *R207K^b^*^6^ showed little difference to the WT. The high fluorescence state of *xL215^b^*^6^ and *G216^b^*^6^ suggested they were blocked in State 1 whereas *R207K^b^*^6^ showed a decreased F_M_’ upon dark anaerobic adaptation, *i.e.* transited to State 2 in response to the reduction of the PQ pool (see Supplementary Figure S3 for the kinetics of F_M_’ decrease corresponding to the false color images of Figure 2). All strains had a similar F_V_/F_M_, showing the maximal quantum yield of PSII was not affected. F_V_/F_M_ was even larger for mutants *stt7-1*, *xL215^b^*^6^ and *G216^b^*^6^, consistently with a larger PSII antenna size expected for State 1.

**Figure 2.**
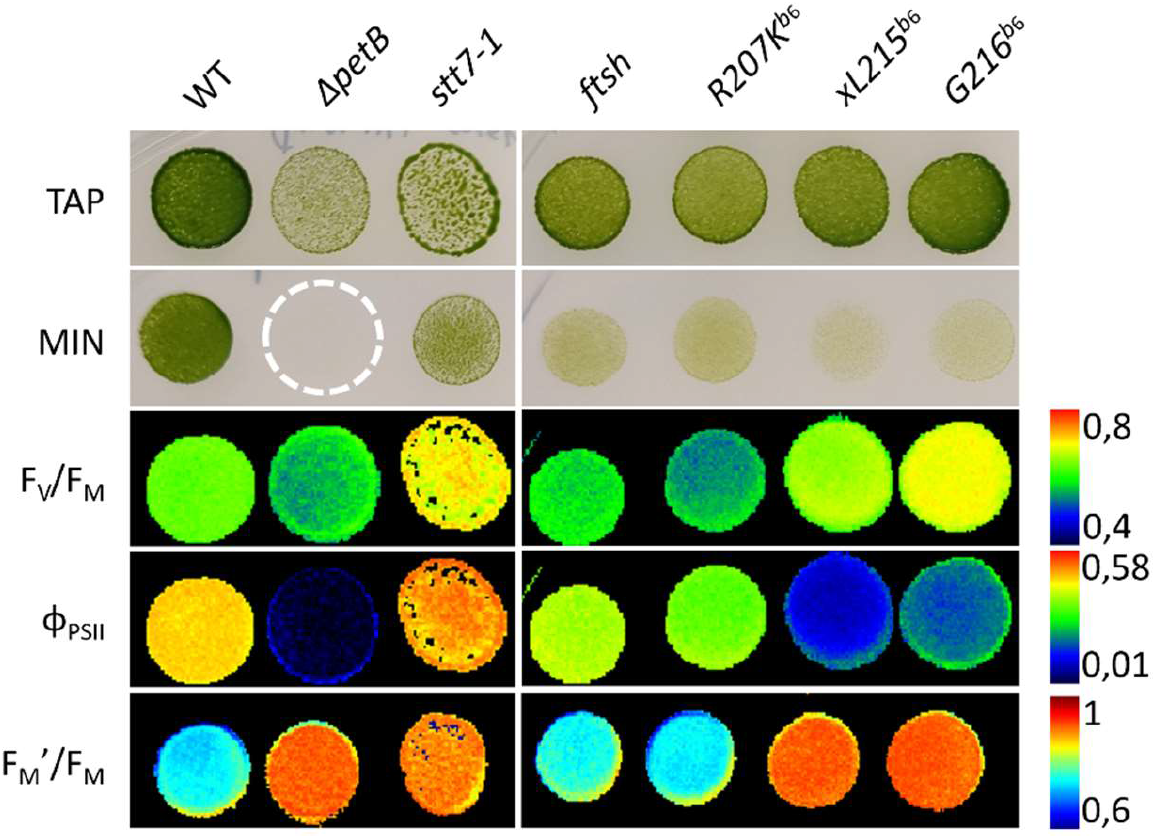
Mutant strains generated for this study and physiological characterization of their photosynthetic properties: heterotrophic growth on tris-acetate-phosphate (TAP, at 5-10 μmol_photons_.m^-2^.s^-1^) and photoautotrophic growth on mineral media (MIN, at 20 μmol_photons_.m^-2^.s^-1^). Maximum quantum efficiency of PSII F_V_/F_M_ = (F_M_-F_0_) / F_M_, quantum efficiency of PSII in the light ϕ_PSII_ = (F_M_’-F) / F_M_ were measured under aerobic conditions using a Chl a fluorescence imaging camera. Fluorescence changes induced by state transitions F_M_’/ F_M_ were measured on aerobic F_M_ compared to F_M_’ quenched after 30 minutes of dark anaerobic adaptation (N_2_ flushed on the plates). For numerical values, on the false color images see color bars on the right. For the whole kinetics of dark anaerobic adaptation extracted from the time-resolved imaging fluorimeter, see Supplementary Figure S3.

The increase of PSI antenna size in State 2 was detected using chlorophyll fluorescence emission spectra at 77K (see Supplementary Figure S4). State 2 was observed for the WT, *ftsh* and *R207K^b^*^6^ but absent in *stt7-1*, *xL215^b^*^6^ and *G216^b^*^6^ supporting a block of state transitions in these strains. Cytochrome *b*_6_*f* complex turnover (Q-cycle) was measured *in vivo* using flash-induced absorbance changes: electrochromic shift (ECS) and cyt *b* and *f* oxidation-reduction kinetics (see Supplementary Figure S5 and Text S1). These measurements showed that cyt *b*_6_*f* complexes accumulated to WT amounts in *R207K^b^*^6^, *xL215^b^*^6^ and *G216^b^*^6^. Although the Q-cycle of *R207K^b^*^6^ was not affected, *xL215^b^*^6^ and *G216^b^*^6^ were blocked in the plastoquinone reduction site Q_i_.

### Truncation or elongation of cyt *b*_6_ impedes the binding of heme *c*_i_

Accumulation of various protein subunits were detected by total protein extraction from a culture grown in TAP in low light (5-10 μmol_photons_.m^-2^.s^-1^), SDS-PAGE and immunodetection using specific antibodies (Figure 3). While *ΔpetB* , was devoid of cytochrome *b*_6_ subunit ^12^, and *stt7-1* lacked the STT7 kinase (Figure 3A), *ftsh* and *R207K^b^*^6^ accumulated these proteins to WT levels. *xL215^b^*^6^ and *G216^b^*^6^ mutants also accumulated normal levels of STT7 and cyt *b_6_*. Truncated cyt *b*_6_ (*xL215^b^*^6^) ran slightly faster on the gel than the elongated form (*G216^b^*^6^) but for both, the bands were much less diffuse than in all other samples (Figure 3C). Similar changes in polypeptide mobility were reported for mutants affected for heme binding ^13^. Heme staining (TMBZ, tetramethylbenzamidine peroxidase activity) showed absence of cyt *b*_6_ and a 90 % reduction of cyt *f* in *ΔpetB* while cyt *f* was detected in all other strains (Figure 3D). Only *xL215^b^*^6^ and *G216^b^*^6^ showed absence of heme staining even though the *b_6_* polypeptide was detected (Figure 3C). Our conclusion is that heme *c*_i_ does not bind covalently to the truncated or elongated versions of the cyt *b*_6_ subunit, and that modified complexes accumulate when FTSH is inactivated.

**Figure 3.**
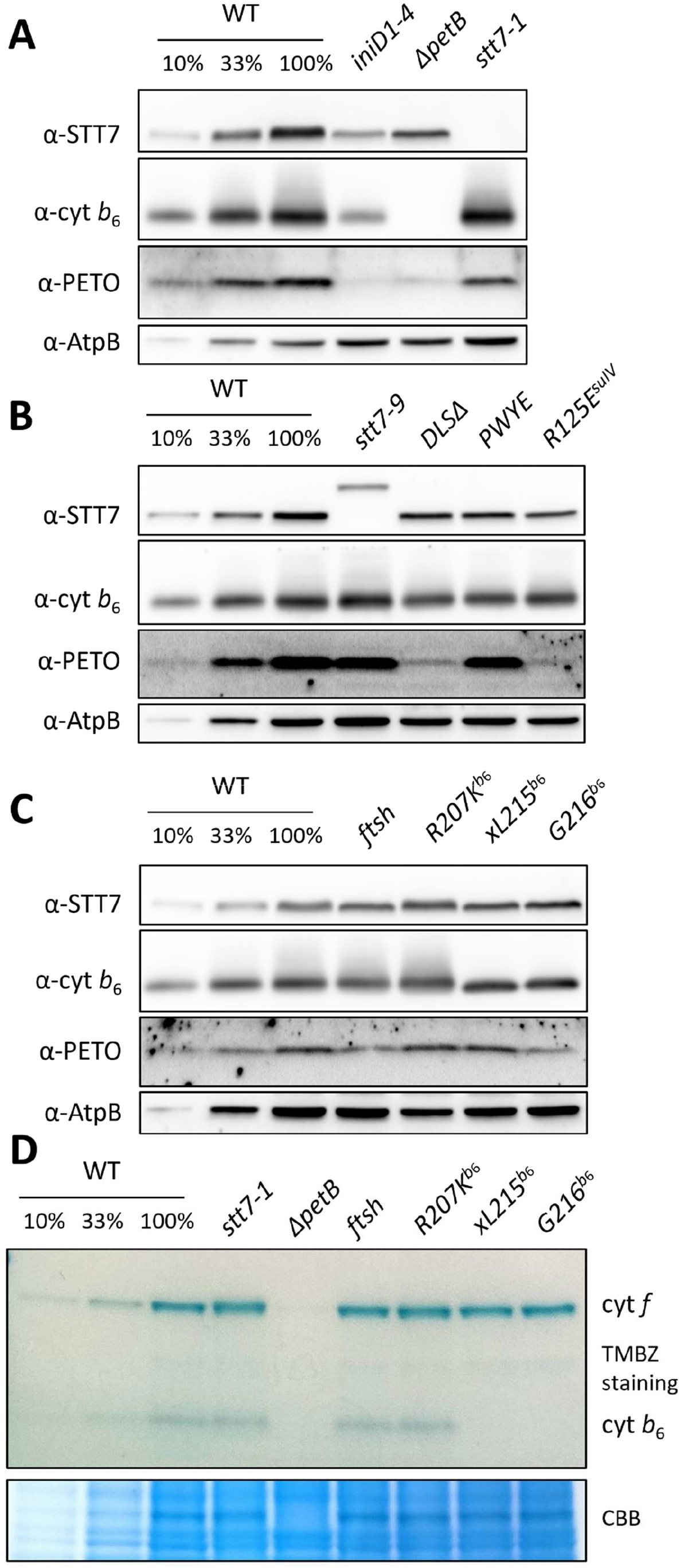
Accumulation of protein subunits in various control strains and mutants affected for state transitions. Polypeptides, isolated from heterotrophically grown cells, were separated by SDS-PAGE and detected with specific antibodies raised against STT7, cytochrome *b*_6_ and PETO. AtpB was used as loading control (see panels A, B and C). Strain *iniD1-4* accumulates reduced amounts of cyt *b*_6_*f* complex and the knock-out mutant *ΔpetB* is completely devoid of the complex. The *stt7-1* mutant is devoid of the STT7 protein kinase while in *stt7-9* a different genetic lesion produces smaller amounts of a higher molecular mass of STT7. *PWYE* and *R125E^suIV^* are site-directed mutants of *petD*, *DLSΔ* is *petD* fused with *petL*. The reference strain *ftsh* is a transformant of *ftsh1-1* with a plasmid carrying non-mutated *petB* while *R207K^b6^*, *xL215 ^b6^* and *G216 ^b6^* are *petB* mutants. Panel D shows a gel with heme staining (TMBZ peroxidase activity). Cyt *f* and cyt *b*_6_ do not accumulate in *ΔpetB*. While cyt *b*_6_ subunit is present (panel C), *xL215 ^b6^* and *G216 ^b6^* show an absence of heme staining of this polypeptide (absence of heme *c*_i_). Cyt *f* staining is equal to WT. Coomassie blue colloidal staining was used as loading control.

*stt7-9* accumulated less of a higher molecular version of STT7, as previously reported ^14^ and attributed to the expression of a putative STT7-Arg fusion. PETO (also referred as suV) is a subunit loosely bound to cyt *b_6_f* complex ^15, 16^. PETO was reduced in *ΔpetB*, *iniD1-4* ^17^*, DLSΔ* ^18^ and *R125E^suIV^* ^9^ (see Figure 3B), suggesting a loss of interaction of PETO with suIV in these strains.

### STT7 phosphorylation in response to reducing conditions

STT7 phosphorylation state was analyzed by polypeptide separation on PhosTag-SDS-PAGE and immunodetection with STT7 antisera. Oxidizing conditions (ox) favoring State 1 were obtained after 1 hour preillumination in the presence of DCMU under oxic conditions; and reducing conditions (red) were reached 10 min after addition of 5 μM FCCP in the dark. Sampling of the proteins was done by fast quenching of cultures in cold acetone. When Phos- Tag is incorporated to denaturing gels, the dephosphorylated polypeptide runs faster than the phosphorylated polypeptide. Here in Figure 4, we denoted D−, P−, and PP− the three bands recognized by the STT7-antiserum. In *stt7-1*, used as a negative control, neither in oxidizing nor in reducing conditions could we detect non-specific bands (see the two rightmost lanes in Figure 4A). In all other strains placed in oxidizing conditions, STT7 was detected as a single band, suggesting that the protein kinase accumulated under its (D−) dephosphorylated form. After the WT was shifted to reducing conditions, this single band disappeared, the migration of STT7 was significantly retarded and split into (P−) phosphorylated and (PP−) highly phosphorylated forms. The heterogeneity in STT7 phosphorylation levels was always, within experimental accuracy, in favor of the highly (PP−) phosphorylated forms (for example compare 4A, 4B and 4C). In the *DLSΔ* mutant, in which *petD* fusion to *petL* gives an extra helix to suIV, destabilizes STT7 binding and diminishes state transitions, P− is accumulated at the expense of PP−. Similarly, in *iniD1*-*4*, attenuated in cyt *b*_6_*f* complex, P− dominates over PP−. In cyt *b*_6_*f* -less mutant *ΔpetB*, only P− is found, as it is in mutants accumulating cyt *b*_6_*f* to WT levels, but blocked for state transitions (*PWYE*, *R125E^suIV^*, *xL215^b^*^6^, *G216^b^*^6^). *ftsh* and *R207K^b^*^6^ showed a distribution of STT7 between the three D−, P− and PP− forms with less immunoreaction. This attenuation of bands in PhosTag-PAGE was inconsistent with the similar levels of STT7 detected in Figure 3C. Therefore, we decided to address the time-dependence of the phosphorylation pattern.

**Figure 4.**
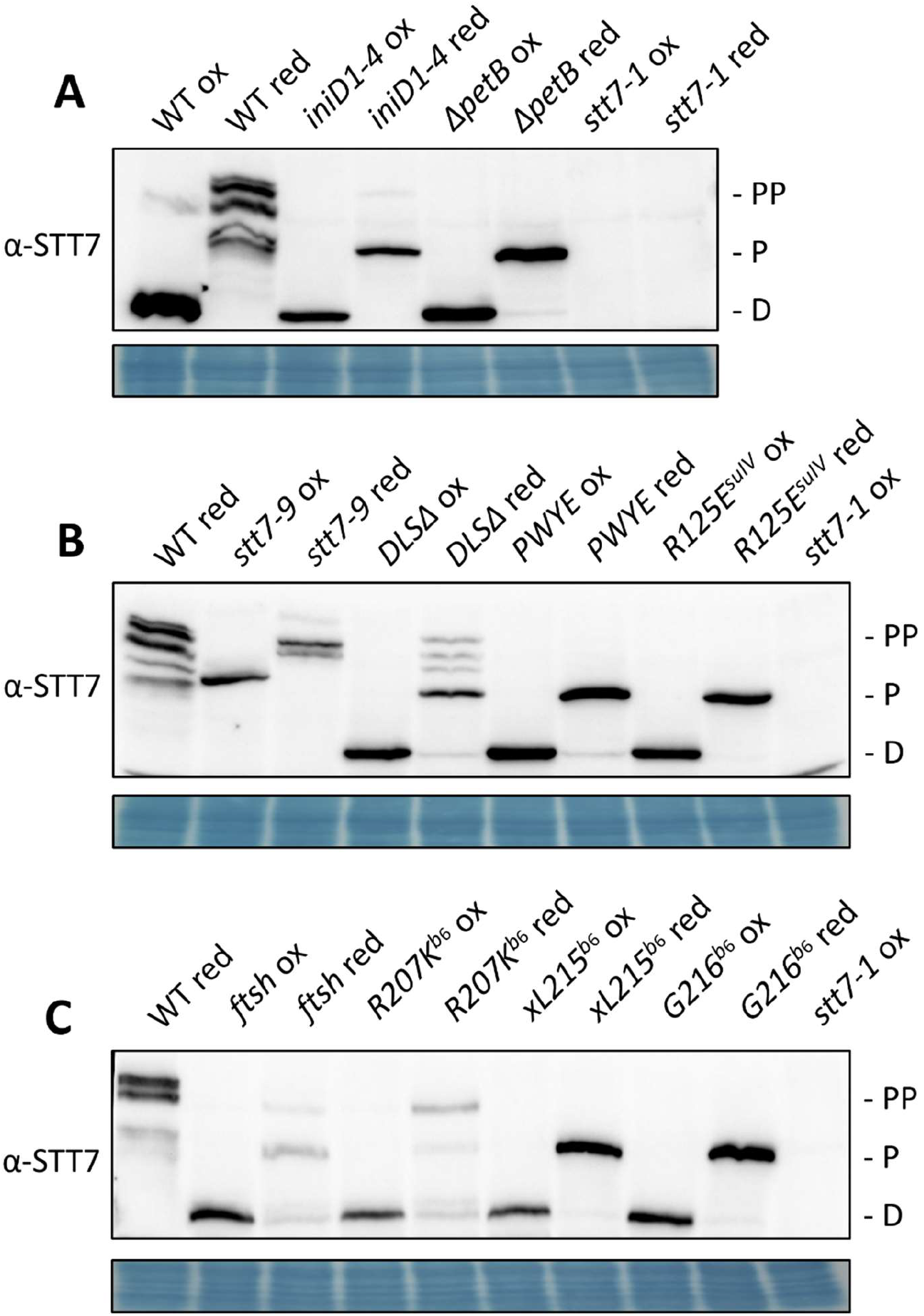
Dynamic changes in STT7 phosphorylation revealed by Phos-tag SDS-PAGE. Phos-tag delays the migration of the phosphorylated forms of polypeptides. Prior to the experiment, oxidizing conditions (ox-label) correspond to a 60 min preillumination in the presence of DCMU (State 1 in the WT). Incubation with 5µM FCCP for 10 min induces the dark reduction (red-label) of the PQ pool (State 2 in the WT). STT7 was detected with a specific antibody. While in all oxidizing conditions (ox), STT7 is dephosphorylated (-D), reducing conditions induced a variety of phosphorylated-forms, -P and -PP, accumulated to various levels. Striking differences are observed between mutant strains. When transferred to reducing conditions, all mutants blocked in State 1 accumulate STT7 under its P-form.

### Kinetics of STT7 phosphorylation and state transition

As in the experiments described above, cells were treated with DCMU and incubated for 1 h under light-oxic conditions (ox). At time *t* = 0 these cultures were transferred to the dark, and reducing conditions (red) were induced by addition of 5 μM of the membrane uncoupler FCCP (Figure 5). Proteins were sampled by fast quenching in cold acetone at various delays after the induction. Protein samples were loaded on PhosTag-PAGE and reacted against STT7 antiserum or on standard PAGE and immunoblotted with an anti-phosphothreonin (α-^p^T) antibody. In the WT (Figure 5A), in response to reducing conditions, and within the time-resolution of the sampling (2-5 min) the D−form of STT7 disappeared and the PP−form accumulated. At the same time, P13 and P17 were phosphorylated. At *t* ≥ 10min, the PP−form steadily decreased while the P−form increased. However, P13/P17 stayed phosphorylated during the course of the kinetics.

**Figure 5.**
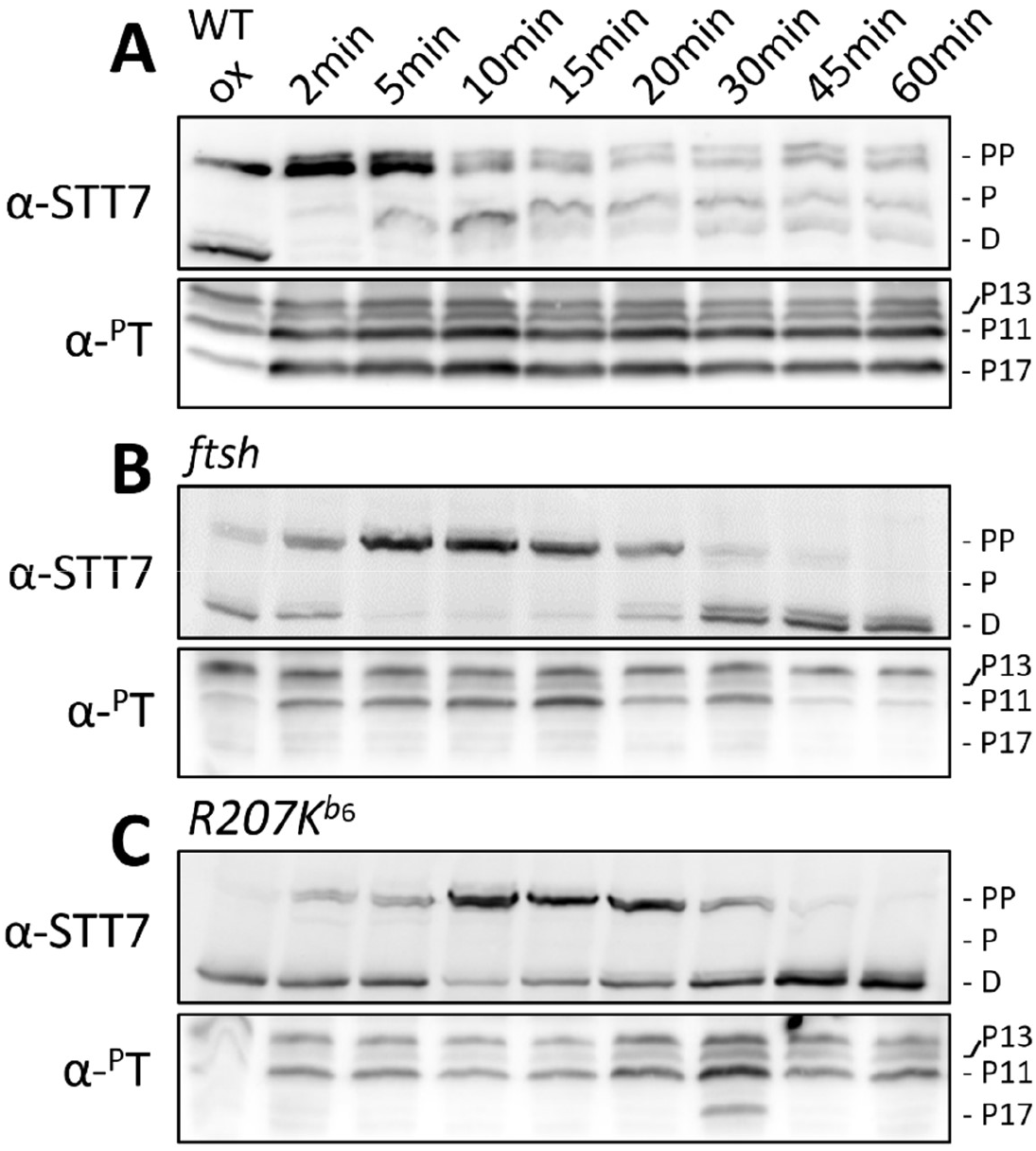
Phosphorylation kinetics of thylakoid polypeptides induced by the reduction of the PQ pool in WT (A), *ftsh* (B) and *R207K^b^*^6^ (C). Prior to the experiment, oxidizing conditions (ox) correspond to a 60 min preillumination in the presence of DCMU. At *t* = 0, the sample was placed in the dark and FCCP was added. Time resolution is limited to 2 min due to protein extraction. Phosphorylation of STT7 was detected with a specific antibody after Tris-acetate Phos-tag SDS-PAGE. Various forms of STT7 were detected: dephosphorylated (-D), mildly phosphorylated (-P) and highly phosphorylated (-PP). Phosphorylation of light harvesting complexes polypeptides was detected after Bis-Tris SDS-PAGE using an anti phosphothreonine antibody (α-^P^T). Phosphorylation of polypeptides P13 and P17 are specific for State 2.

In *ftsh* (Figure 5B), PP−formation was delayed, showing a maximal accumulation at 5 ≤ *t* ≤ 15 min. Phosphorylation of P13/P17 peaked at *t* = 15min. In *R207K^b^*^6^ (Figure 5C), PP−formation was further delayed to 10 ≤ *t* ≤ 20 min while the P−form was still detected in a non-negligible amount. P13/P17 phosphorylation peaked at *t* = 30 min. At *t* ≥ 30 min, PP− and P−form were destabilized in both *ftsh* and *R207K^b^*^6^, and STT7 returned to its D−form.

### Identification of phosphorylated sites on STT7

The *Chlamydomonas reinhardtii* (*Cr*) serine/threonine protein kinase STT7 fits the canonical protein kinase fold with a small N-terminal lobe of antiparallel β-strands and a C-terminal lobe rich in α-helices. The hinge between lobes as well as the catalytic loop (hrD_302_xKxxN_307_) are followed by the metal-binding motif (D_321_LG). Similarly as for the cyclin-dependent protein kinase 2 (CDK2), *Micromonas* species (*Ms*) and *Arabidopsis thaliana* (*At*), CrSTT7 has an ATP-binding site involving K_172_*^Cr^*^STT7^, N_307_*^Cr^*^STT7^ and D_321_*^Cr^*^STT7^. These residues are underlined in Figure 7, together with other residues shaping the ATP-binding site in *Micromonas*. Compared to *Ms*STT7 and *At*STN7, *Cr*STT7 has two insertions, in grey in the figure, between the α-helices 8 and 9 and between the β-strands 5 and 6. Although none of these affect the protein fold, the sequence alignment is rather wobbly in these regions. The former insertion likely occurred between S_205_*^Cr^*^STN7^ and S_225_*^Cr^*^STN7^ in Chlamydomonas, Arabidopsis having two consecutive serines in this region S_202_S_203_*^At^*^STN7^. Previous phosphoproteomics studies showed phosphorylation of S_533_*^Cr^*^STN7 19^ and S_526_*^At^*^STN7^ (Reiland 2009) (Trotta 2016) (see Figure 7A). These Ser or Thr residues do not align between Chlamydomonas and Arabidopsis, and not only are they dispensable for state transitions ^20^, but also is the whole C-terminal extension, absent from Micromonas. The recombinant STT7 kinase catalytic domain produced from *E. coli* and reconstituted with cyt *b*_6_*f* complex showed ATP hydrolysis and STT7 autophosphorylation ^9^. We therefore used our purified truncated version of the kinase to sharpen the detection of phosphorylated peptides by mass spectrometry, and to be more specific to the catalytic site of STT7. Proteomics (see Supplementary Excel file) covered 96% of the STT7 protein sequence (and 100% of the Ser or Thr residues). We detected several peptides identifying 6 different phosphorylation sites, including 4 previously unknown sites in the catalytic domain of STT7 (see Supplementary Text S2 for sequence alignment): (a) T_152_, (b) T_166_ or S_167_, (c) T_200_ or S_205_, (d) S_295_, (e) Y_465_, (f) T_489_, S_491_, or S_495_.

**Figure 7.**
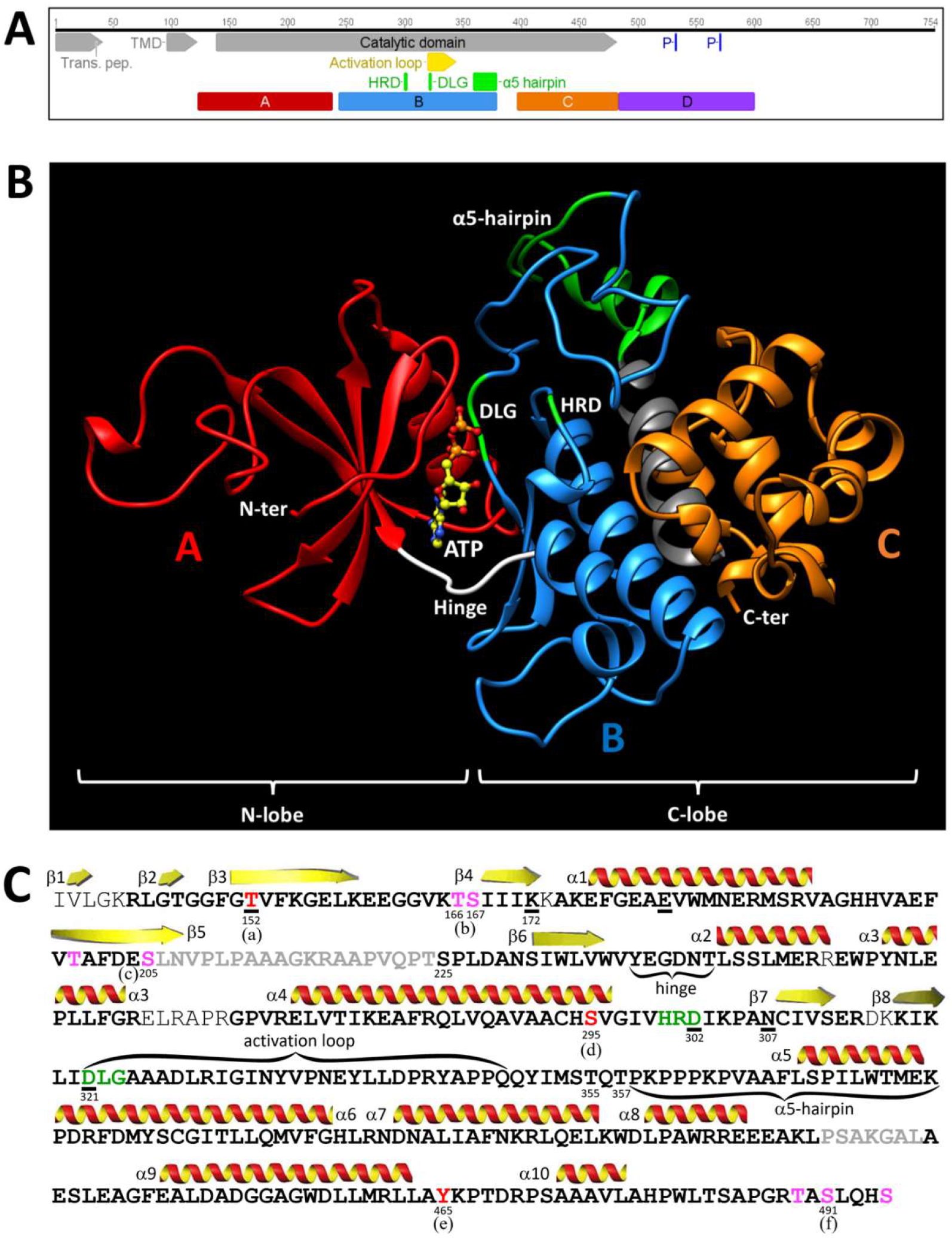
Structural features of the STT7 protein kinase showing the transmembrane domain (TMD), the kinase catalytic domain (panel A) and the previously reported phosphorylatable residues in the C-terminal extension ^19, 20, 30^. 3D model of the truncated CrSTT7 kinase domain on the crystal structure of Micromonas STT7 (panel B). Produced as a recombinant protein in *E. coli*, the A (N-lobe) and B-C (C-lobe) domains are sufficient for kinase activity ^9^. Phosphoproteomics (panel C) covered 96% of the sequence (bold), 6 phosphorylation sites were evidenced in this work, and noted (a-f). These phosphorylatable residues are coloured in red (only one per phosphopeptide) or pink (>1 per phosphopeptide). Canonical protein kinase catalytic loop (HRD) and metal-binding motif (DLG) are coloured in green. The residues forming the ATP-binding site are underlined.

## Discussion

### The C-term of cyt *b*_6_ straps the complex subunits together

The chloroplast cytochrome *b*_6_*f* complex is involved in photosynthetic electron transport while its mitochondrial counterpart, the cytochrome *bc*_1_ complex is involved in respiration. Apart from the difference between cyt *c*_1_ and cyt *f*, which play very similar roles in terms of electron transport, the most significant genetic difference is that the bacterial and mitochondrial *petB* gene is split into *petB* and *petD* in chloroplast or cyanobacterial version of the *b*-type cytochrome ^21^. The *petB-petD* split co-occurs with the presence of C35*^b^*^6^, forming the thioether bond to the *c*_i_ heme. Therefore, the fragmentation of the core subunits of the complex was thought ^10^ to accommodate heme *c*_i_ insertion ^22^. Here we show that the truncation (*xL215^b^*^6^) or elongation (*G216^b^*^6^) of the cyt *b*_6_ does not allow for cyt *b*_6_*f* accumulation. If cyt *b*_6_*f* accumulation is favored when FTSH is inactivated, heme *c*_i_ does not bind covalently to C35*^b^*^6^. We therefore conclude that the salt-bridge formed between the C-term of cyt *b*_6_ and suIV straps these two subunits together. When loosened (*G216^b^*^6^) or fastened (*xL215^b^*^6^), the strap fails to stabilize the CCB2-4/CCB3/cyt *b*_6_ transient heme *c*_i_ ligation complex ^23^. The final step of assembly of the complex being aborted, modified complexes are degraded by FTSH.

### The stromal side of the cyt *b*_6_*f* complex is a protein docking site for STT7, PETO and other proteins

For the functional study of the modified cyt *b*_6_*f* complexes, we used *ftsh1-1* as a host for the genetic engineering ^24^. We found in *G216^b^*^6^ and *xL215^b^*^6^, that cyt *b*_6_*f* subunits accumulated to WT levels, including PETO (also referred to as suV) ^3^. In contrast, PETO accumulation is reduced *DLSΔ* and *R125E^suIV^*. We take this as a support to the crosslinking data and model of interaction between PETO and suIV ^16^, and suggest that unbound PETO gets degraded by FTSH. Although PETO is phosphorylated in State 2 ^15^, the presence or absence of PETO does not have any influence on state transitions ^25^. This algal-specific protein may rather stabilize cyt *b*_6_*f* supercomplexes under anaerobic conditions ^25^. In cyanobacteria, R125^suIV^ binds to PETP ^26^ instead of PETO ^25^ or STT7 ^9^. As for Chlamydomonas PETO, cyanobacterial PETP may regulate the dynamic interaction of cyt *b*_6_*f* complex with other protein complexes ^27^. In the spinach cyt *b*_6_*f* complex, the thylakoid soluble phosphoprotein (TSP9) is docked to the cyt *b*_6_ – suIV stromal interface ^28^. The binding / release mechanism proposed below for STT7 may work similarly for PETP and TSP9, providing a regulatory control on the signaling or metabolic pathways involving PETP and TSP9.

#### A phosphorylation model for state transitions

In Figure 4A we showed a phosphorylation of STT7, from the dephosphorylated form D− to the mildly phosphorylated P−form, that occurred in the absence of the cyt *b*_6_*f* complex. This is represented in Figure 6 as steps ① and ②, independent from cyt *b*_6_*f* complex. However, it does not indicate that the STT7 P−form ② is required for binding to the cyt ③. Although not involved in the phosphorylation step from ① to ②, cyt *b*_6_*f* may bind the D−form ① (see dotted line) as well as the P−form ② (solid line). Despite cyt *b*_6_*f* being 20 times more abundant than the protein kinase ^7^, we showed here that a ∼10-fold decrease in the amount of cyt *b*_6_*f* becomes limiting for STT7 phosphorylation (see *iniD1*-*4* lane in Figure 4A).

**Figure 6.**
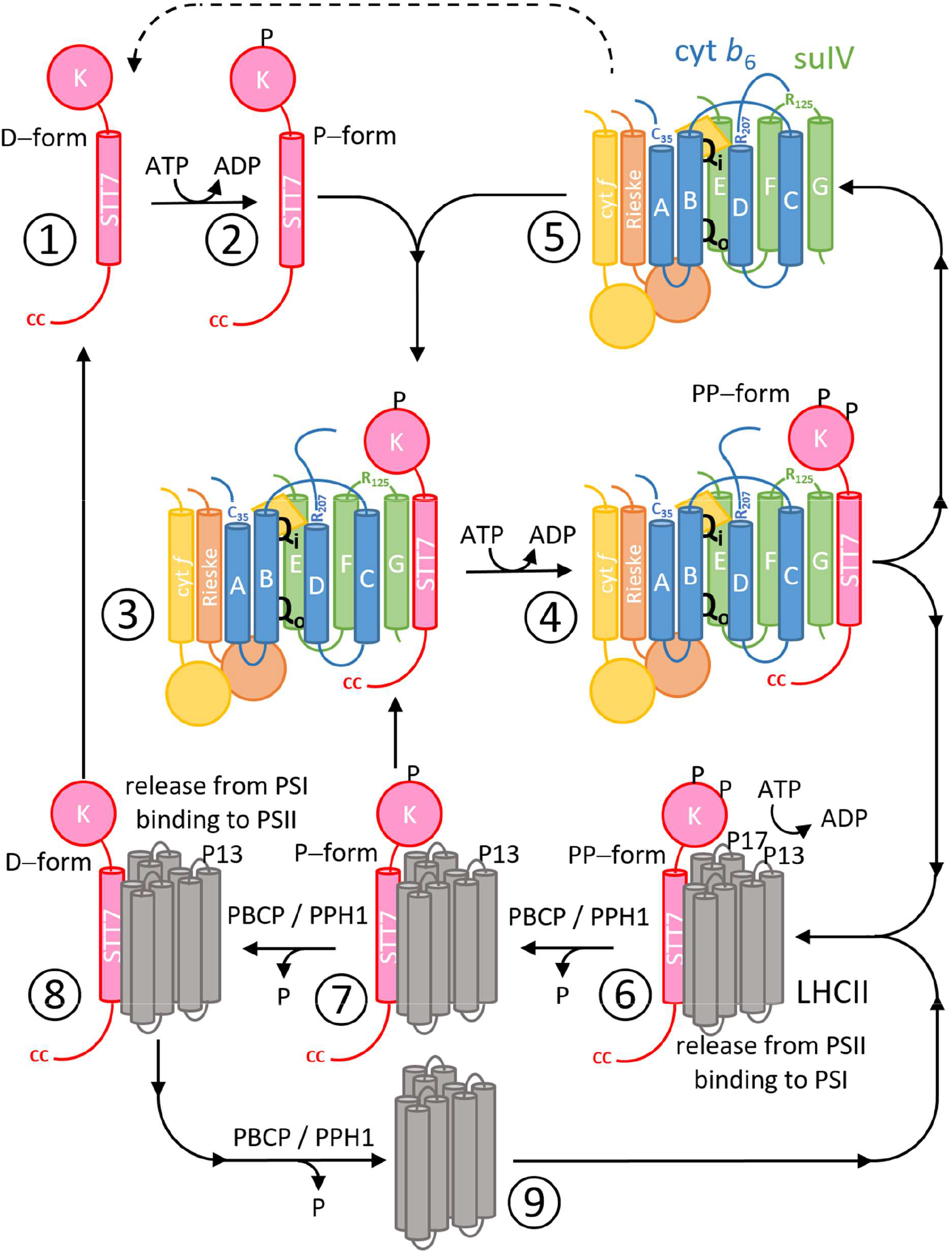
Model for cyt *b*_6_*f* complex – STT7 protein kinase interaction, phosphorylations and state transitions. (see text for details). The first phosphorylation step ① to ② is independent from cyt *b*_6_*f* complex (see *ΔpetB* in Figure 4A). STT7 binds to the F-G interface of subunit IV and interacts directly with R125^suIV^, steps ⑤ to ③. The cytochrome is required for the formation of the highly phosphorylated form of STT7, step ③ to ④ and the phosphorylation of P13 and P17 of light harvesting complexes II. PBCP and PPH1 phosphatases dephosphorylate LHCII, steps ⑥ to ⑨.

We have shown a direct interaction between cytochrome *b*_6_*f* complex suIV and the STT7 protein kinase, which induces STT7 autophosphorylation when ATP is added ^9^. This is represented in Figure 6 as central steps ③ to ④. STT7 is always found attached to the complex ^7^, meaning that the reaction steps ③ and ④ are the most stable. It suggests that the fraction of STT7 diffusing freely in the membrane is very small or that the STT7 kinase catalytic domain just flips from suIV to the neighbouring LHCII complexes.

The activation of STT7 through the highly phosphorylated PP−form in step ④, is necessary for the phosphorylation of LHCII, represented here as step ⑥. In the WT strain, the full phosphorylation of LHCII is reached in less than 2 min of incubation under reducing conditions, *i.e.* much faster than the kinetics of state transitions. It shows that the limiting step for state transitions is not the phosphorylation of LHCII but the migration of LHCII from PSII to PSI. When the formation of the highly phosphorylated forms of STT7 ④ is slowed down, like in the *R207K^b^*^6^ substitution mutant, the sampling of the reaction steps allows for a better time-resolution of the phosphorylation kinetics. In Figure 5C, the phosphorylation of the P13 polypeptide seems to occur before P17, and P13 remains phosphorylated to the detriment of P17, see step ⑦. P13 and P17 appear phosphorylated at the expense of the STT7 PP−form, which returns to the P− or to the D−forms, the latter being strongly favoured at 60 minutes in the *R207K^b^*^6^ mutant. It suggests the PBCP / PPH1 phosphatases have a lesser affinity for P13, step ⑧ to ⑨ (see Figure 5C).

At this stage, it is still unclear how many phosphorylation reactions occur in the transient STT7-LHCII complex at step ⑥. Activated STT7 PP−form may hydrolyse ATP to phosphorylate several LHCII complexes. It is also possible that STT7 PP−form transfers a phosphate to LHCII and requires cyt *b*_6_*f* reactivation to reset to the PP−form.

### STT7 phosphorylated residues

Protein kinases activation occurs via autophosphorylation ^29^. Previous studies identified some phosphorylated STT/N7 peptides from the C-terminal region of the protein ^14, 19, 30^. The complete protein coverage of our proteomics study showed that STT7 gets phosphorylated at 6 different sites in the catalytic domain. This is consistent with the multiple, sometimes split bands we detect using Phos-Tag PAGE. The six different phosphopeptides we reported here were:

a. Phosphopeptide containing T_152_*^Cr^*^STT7^ is also found phosphorylated in Micromonas ^31^ (T_185_*^Mc^*^STT^^7^) but is not conserved in Arabidopsis.
b. Phosphopeptide containing non-conserved T_166_ and S_167_*^Cr^*^STT7^.
c. Phosphopeptide containing non-conserved T_200_*^Cr^*^STT7^, and S_205_*^Cr^*^STT7^ that is conserved in Arabidopsis (aligned with S_202 or 203_*^At^*^STN7^), but not in Micromonas.
d. Phosphopeptide containing S_295_*^Cr^*^STT7^ conserved in Arabidopsis (S_272_*^At^*^STN7^) and close to S_305_*^Mc^*^STT7^ in Micromonas.
e. Phosphopeptide containing Y_465_*^Cr^*^STT7^ conserved in Arabidopsis (Y_435_*^At^*^STN7^), but non-conserved in Micromonas.
f. Phosphopeptide containing T_489_, S_491_*^Cr^*^STT7^, and S_495_. This phosphopeptide was also reported for Chlamydomonas ^14^. Only S_491_*^Cr^*^STT7^ is conserved in Arabidopsis (S_461_*^At^*^STN7^) but *Mc*STT7 terminates earlier, shortly after the α-10 helix.

We confirmed two sites identified earlier (a) ^31^ and (f) ^14^. Half of the phosphorylated sites are found in the ATP-binding pocket (a), (b) and (d), near the underlined residues in Figure 7C. Similarly as proposed for site (a) ^31^, we propose that these phosphorylations are induced by ATP hydrolysis. The other identified sites (c), (e), and (f), are more distant from the ATP-binding site and may get phosphorylated by another kinase. Only the new phosphorylation site (d) at S_295_*^Cr^*^STT7^ is conserved in Arabidopsis (S_272_*^At^*^STN7^), only 4 residues downstream of Micromonas S_305_*^Mc^*^STT7^.

In human PAK2 or AKT1 kinases, auto-phosphorylation occurs in the activation loop, inducing a conformational change and activation of the enzyme^29, 32^. Plant STT/N7 kinases do not have such a phosphorylatable residue in this protein domain. Instead, they have an extra α-5 hairpin domain. Here we did not find phosphorylation on T_355_ or T_357_*^Cr^*^STT7^, and we propose that protein phosphorylation at S *^Cr^*^STT7^, at the C-term end of the largest α-4 helix, just upstream of the catalytic loop (hrD_302_xKxxN_307_) is the trigger for protein conformational change in STT/N7 kinases.

### Role of the quinone binding sites Q_o_ and Q_i_

In support to previous work ^4, 5^, we confirm here that the quinone binding site Q_o_ plays a pivotal role in state transitions: Figure 4B shows that STT7 is blocked under its P−form in the PWYE mutant. In accordance with STT7 being blocked in the P−form in the strain devoid of cyt *b*_6_*f* complex, a functional Q_o_ site, and movement of the Rieske protein ^6^, may be required for the binding of cyt *b*_6_*f* complex to STT7 through its transmembrane helix and luminal extension.

In the cyt *b*_6_*f* complex crystal structure, the C-terminal carboxylate group of cyt *b*_6_ interacts with the guanidinium group of R125^suIV^ ⑤. *In vivo*, this salt-bridge can be broken to allow R125^suIV^ to interact with STT7 ③ and ④ ^9^. A similar protein-protein interaction is found in the α1α2β1β2 complex of deoxyhemoglobin: a salt bridge is formed between the C-terminal of the β1 subunit and the amino group of Lys40 in the α2 subunit. This bond is broken upon oxygen binding to the heme iron (oxyhemoglobin) ^33^. This conformational change accounts for the cooperative binding of the other three oxygen molecules in the complex. In the cytochrome *b*_6_*f* complex, the binding/release of ligands to the heme *c*_i_ iron may induce similar long-distance stereochemical changes. The Q_i_ site can accommodate quinone analogs such as the NQNO inhibitor or heme iron ligands like carbon monoxide (CO) ^34–36^. Both chemicals change heme *c*_i_’s midpoint redox potential. We also found that heme *c*_i_ titrates in two pH-dependent waves in the presence of NQNO, showing an equilibrium between two conformations of the complex ^34^. Taken together, our results suggest that depending on Q_i_ occupancy and conformation of R207 *^b^*^6^ interacting with heme *c*_i_ propionate ^37^, the unpinning of cyt *b*_6_-COOH from R125^suIV^, is a signal for STT7 kinase activation *via* autophosphorylation.

## Materials and Methods

### *Chlamydomonas* strains and growth conditions

The *Chlamydomonas reinhardtii* strains used in this study are listed in Supplementary Table S1. They were maintained under photoheterotrophic conditions on Tris-acetate phosphate (TAP) plates supplemented with 2% agar at 25°C under 5-10 μmol_photons_.m^-2^.s^-1^. Liquid cultures were grown in liquid TAP medium under the same light intensity unless indicated otherwise. Light was provided by “warm white” LEDs. WT t222 (IBPC, Paris), derivated from 137c ^38^, was used as a reference strain in this work.

### Plasmids and *petB* mutants

Plasmid pWBA containing the *aadA* (spectinomycin-resistance) cassette ^39^ downstream of WT *petB* was used for mutagenesis and chloroplast transformation. When transformed in the *ΔpetB* strain, the site-directed mutated strains were noted *R207K, xL215*, and *G216* see Supplementary Figure S1. The *ftsh 1.1* strain ^11^ was then used as the host strain for chloroplast transformation. The control strain, *ftsh* (*ftsh 1.1* transformed with non-mutated pWBA plasmid) and site-directed mutated strains (note the *b*_6_ superscript) *R207K^b^*^6^*, xL215^b^*^6^ (truncation of the cyt *b*_6_ by deleting L215) and *G216^b^*^6^ (elongation of the cyt *b*_6_ by adding a Gly at position 216) were obtained using mutated pWBA plasmids as follows. Plasmid DNA was precipitated on gold particles using the Seashell Technology S550d gold DNA protocol. *ftsh1-1* cells spread on TAP medium with 150 μg.mL^-1^ spectinomycin were transformed by gold particle bombardment. After transformation, cells were left to recover in the dark for up to 24 h and transformants were incubated under low light (5-10 μmol_photons_.m^-2^.s^-1^). Transformants were transferred repeatedly on TAP + spectinomycin medium until homoplasmy was reached.

### Genotyping

DNA extraction was performed using the “Chelex” method. Clones were resuspended in 30 μL H_2_O, 30 μL of ethanol was added and the mix was vortexed for 3s. 200 μL of 5 % Chelex 100 solution (Bio-Rad) was added before 8min incubation at 95 °C. Resin was pelleted by centrifugation and supernatant was used as template for PCR. Genotyping PCRs were performed using WT-specific primers for each mutation. Sensitivity and selectivity of the amplifications were checked using diluted WT DNA extract, see Supplementary Figure S2. Mutations were confirmed by sequencing of the target region of *petB*.

### Chlorophyll fluorescence measurements

Chlorophyll fluorescence was measured on cells grown on plates using a laboratory built system ^40^. Maximum quantum efficiency of PSII [F_V_ / F_M_ = (F_M_ – F_0_) / F_M_], PSII quantum yield [ϕ_PSII_

= (F_M_ – F) / F_M_] under 500 μmol_photons_.m^-2^.s^-1^. PQ pool was oxidized by a 15min dark adaptation before transfer to anaerobic conditions (N_2_ flush) and F_M_^’^ / F_M_ was recorded during a 10 min period in the dark with 200 ms saturating pulses. For more details, see Supplementary Text S1.

### Protein isolation and Immunoblot Analysis

For protein accumulation analysis, cells from cultures at 2-4.10^6^ cells.ml^-1^ were spun down and were lysed in 50mM Tris pH 6.8, 2% SDS and 1X protease inhibitor (P9599, Sigma-Aldrich) at 37 °C for 30 min. Polypeptides were separated on 4-12 % and 12 % denaturing NuPAGE gels with MOPS running buffer. Antisera raised against cytochrome *b_6_*, PETO, β subunit of ATPsynthase AtpB and STT7 kinase as well as anti-phosphothreonine antibodies (Cell Signaling) were used for immunodetection by ECL.

### TMBZ peroxidase activity staining

After polypeptides separation on a 4-12 % NuPAGE gel, heme peroxidase activity was detected by TMBZ (3,3′,5,5′-Tetramethylbenzidine) staining ^41^. The gel was incubated for 1 h in the dark in 6.3 mM TMBZ, 175 mM sodium acetate, 30 % methanol, pH 5 before addition of 30 mM H_2_O_2_. Staining took place at 4 °C in the dark over 72 h. Excess of dye was washed in 175 mM sodium acetate, 30 % isopropanol, pH 5.

### Induction of State 1 and State 2

Cells in exponential phase (2-3.10^6^ cells.ml^-1^) in photoheterotrophic conditions in low light (10 μmol_photons_.m^-2^.s^-1^) were concentrated in fresh medium at 2.10^7^cells.ml^-1^. After 1 h recovery in low light, oxidizing conditions induced State 1 (1 h incubation in the light with 10 μM DCMU). After this pretreatment, State 2 was induced by reducing conditions such as addition of 5 μM membrane uncoupler FCCP (carbonyl cyanide 4-(trifluoromethoxy)phenylhydrazone) in the dark or anaerobic treatment (N_2_ flushing on plates) in the dark.

State transition was monitored using 77 K fluorescence emission spectra. Chlorophyll was excited with a 455 nm LED and fluorescence was selected with a red-colored filter and measured with an Ocean Optics USB2000 CCD spectrometer. *Chlamydomonas reinhardtii* cells were kept frozen at 77 K in liquid nitrogen during measurement.

### PhosTag-PAGE analysis of phosphorylated forms of STT7

To analyze fast and sensitive change of phosphorylation of STT7 kinase, cell cultures were quenched in -20°C acetone at a ratio of 1:4. Precipitated proteins were resuspended in 50 mM Tris pH 6.8, 2 % SDS and desalted using ZebaSpin 7K desalting column (ThermoScientific). Separation of phosphorylated forms was done using Zn-PhosTag (Wako) at 55 μM in Tris-Acetate polyacrylamide gel ^42^. Acrylamide:bis-acrylamide ratio in the resolving part of the gels was 75:1. Proteins were transferred during 16 h in a wet blotting apparatus on 0.45 μm nitrocellulose membrane. Immunoblotting was performed with STT7 antisera and ECL detection.

### Phosphoproteomics analysis

Purified STT7 kinase domain and WT cyt *b*_6_*f* complexes were purified and reconstituted *in vitro* with ATP and polypeptides separated on SDS PAGE ^9^. STT7 bands were cut out, digested by Trypsin, and sent for phosphoproteomics analysis by tandem CID (collision-induced dissociation) analysis on a Q Exactive (Thermoscientific) to determine phosphorylation sites. The One-to-One threading tool of the Protein Homology/analogY Recognition Engine V 2.0 (PHYRE2 ^43^) was used to model the structure of the kinase domain of STT7 based on the crystal structure of the kinase domain of its homolog in Micromonas sp. RCC299 ^31^.

## Acknowledgments

This work was supported by Centre National de la Recherche Scientifique and by the Agence Nationale de la Recherche grant PhotoRegul (ANR-18-CE20-0010-01) and HemeMotion (ANR-22-CE20-0042-01). Daniel Picot is warmly acknowledged for discussions and critical reading of the manuscript. We express our gratitude to Marion Hamon for proteomics analysis.

## Abbreviations

Cyt: cytochrome
stt: state transitions

## Supplementary Information

**Table S1.**
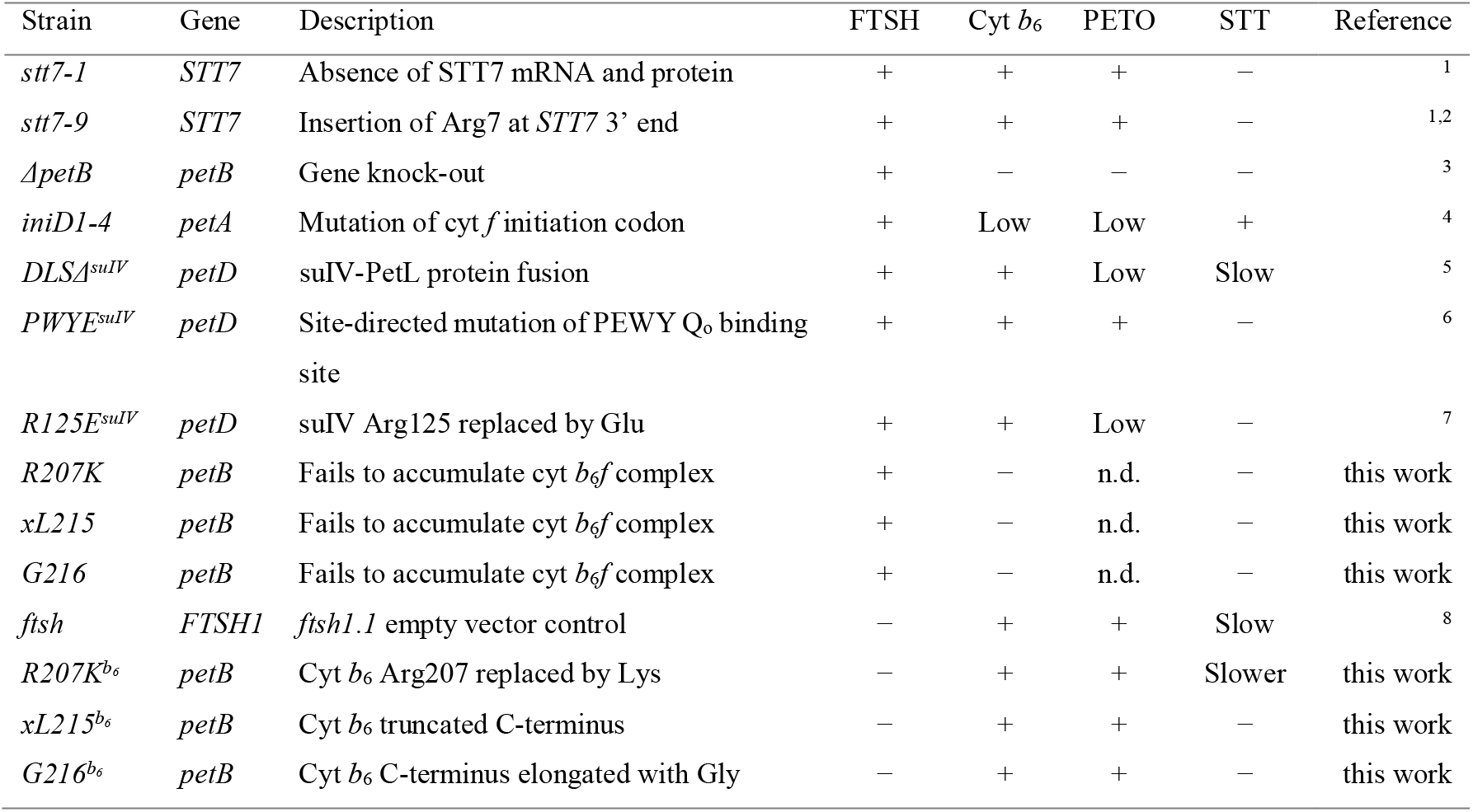
List of mutant strains used in this work, showing the presence (+) or absence (−) of FTSH protease activity, cyt *b*6 and PETO subunits. Most of these strains are impaired for State Transitions (STT).

**Figure S1.**
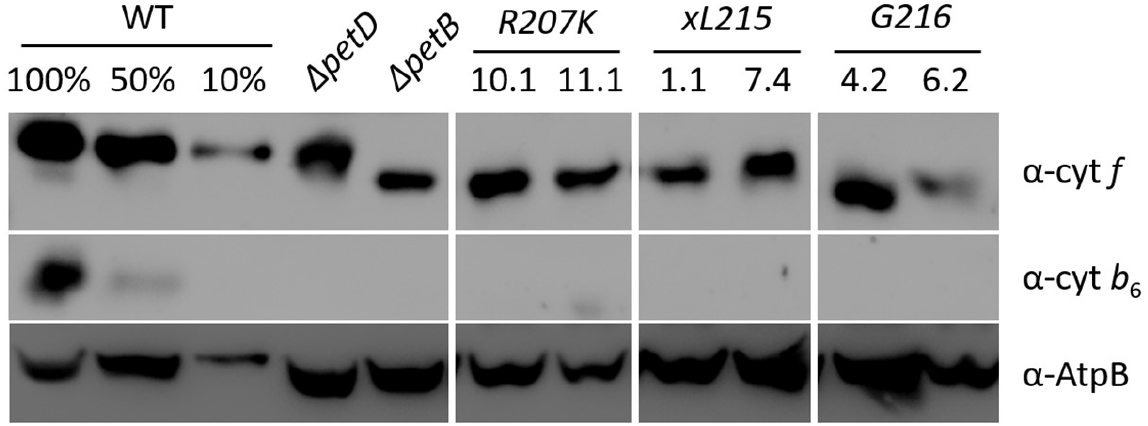
Unsuccessful attempts to obtain modified cyt *b*_6_ when FTSH is active. Transformants of a *ΔPetB* strain were sequenced to verify the presence of the mutated *PetB* genes, but none of these mutants accumulated the cyt *b*_6_ subunit neither did they grow under photoautotrophic conditions.

**Figure S2.**
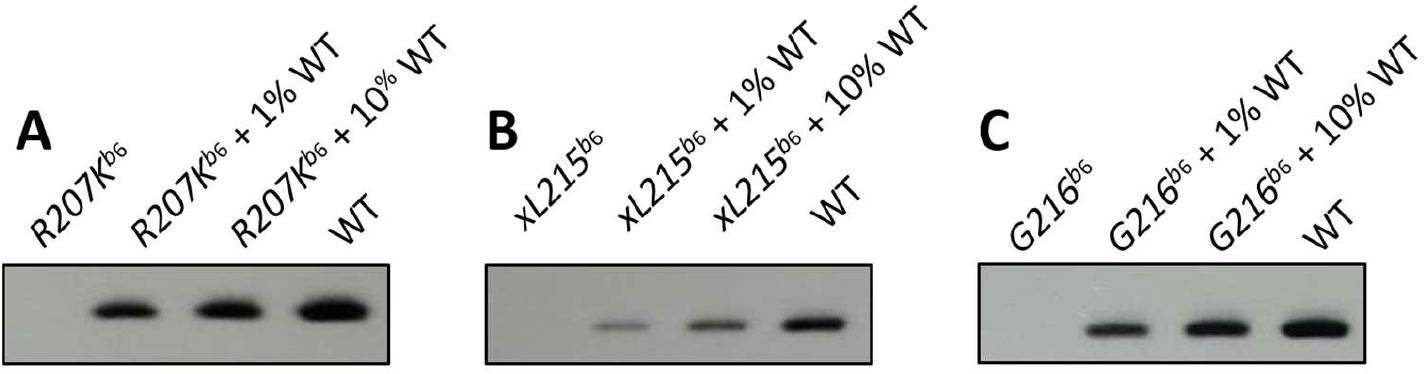
Genetic transformation of *ftsh 1.1* strain provided homoplasmic site directed mutants of the chloroplast *petB* gene. This is shown by WT-specific PCR on DNA extracts of mutants *R207K^b^*^6^(A) *xL215^b^*^6^ (B) and *G216^b^*^6^ (C). 10% and 1% dilutions of WT DNA extract were added to the mutant DNA extract to probe the sensitivity of the PCR.

**Figure S3.**
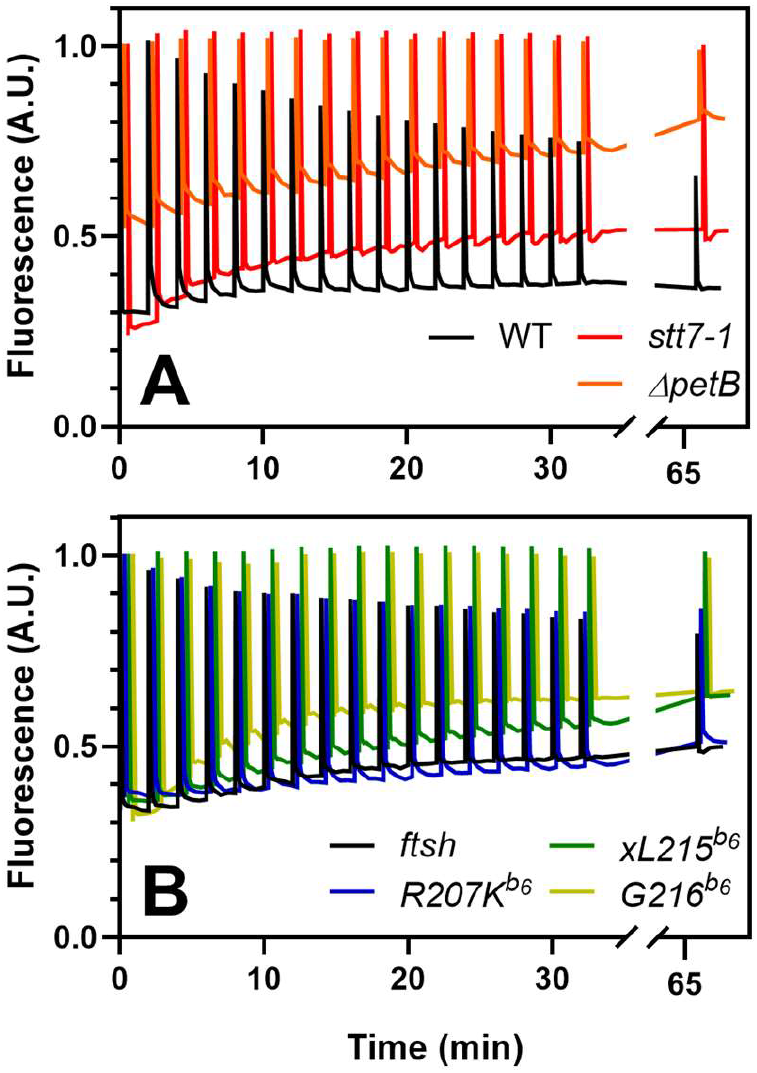
Kinetics of chlorophyll fluorescence quenching upon dark anaerobic adaptation (State 1 to State 2 transition). These data are from the same dataset as in Figure 2 from the main text. Anoxia was induced at *t* = 0 by flushing N_2_ onto the plate. Saturating pulses were used to probe F_M_’. The rise of the steady-state fluorescence F’ is induced by the non-photochemical reduction of the PQ pool (reductive redox poise in anoxic cells). (A) The WT strain shows a quenching of F_M_’ absent from mutants blocked for state transitions (*stt7-1* and *ΔpetB*). (B) While *ftsh* and *R207K^b^*^6^ are similar to the WT, *xL215^b^*^6^ and *G216^b^*^6^ are close to *stt7-1*.

**Figure S4.**
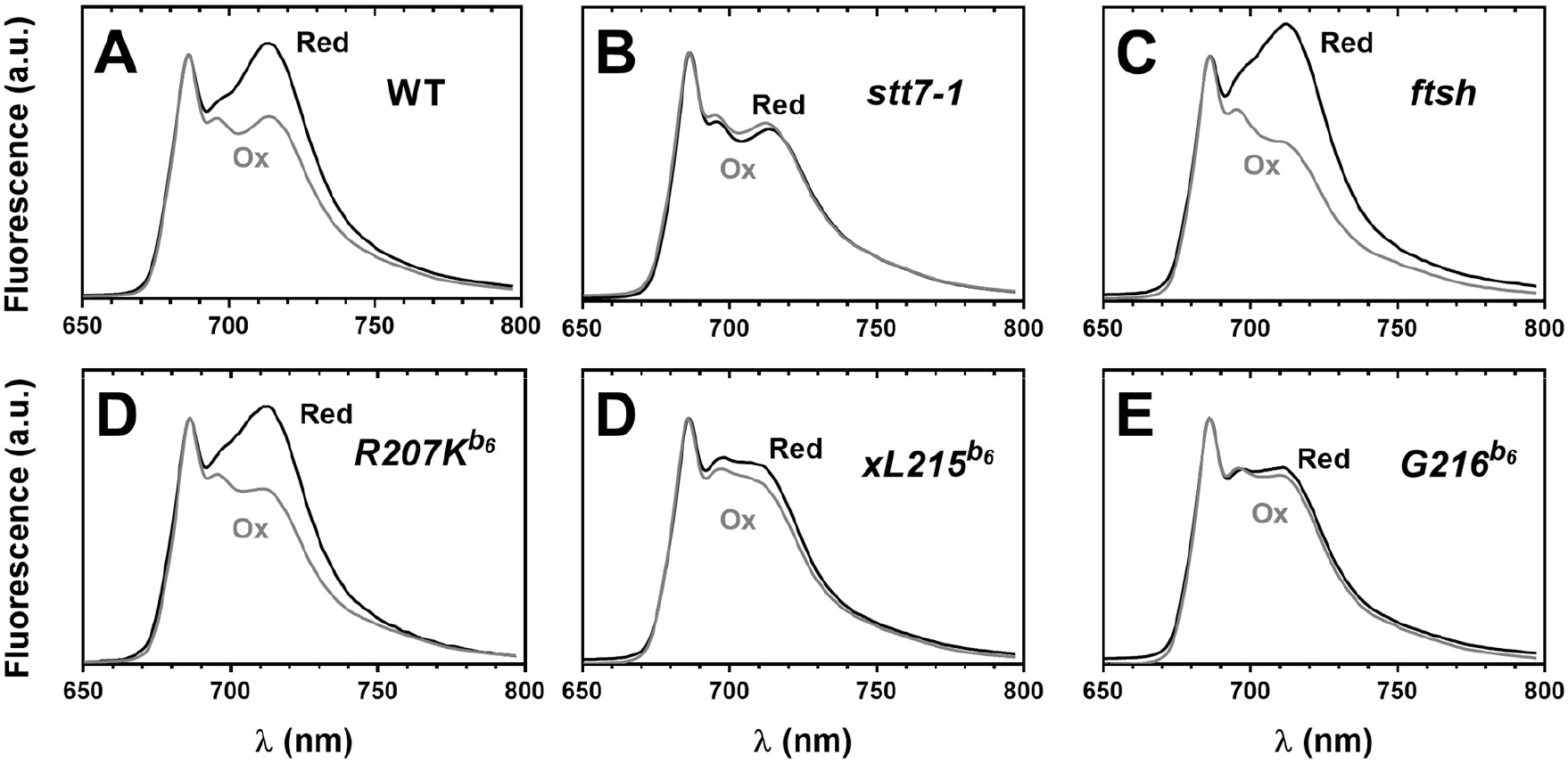
State transitions measured as 77K chlorophyll fluorescence emission spectra. Spectra in oxidizing (Ox) and reducing (Red) conditions were normalized on the PSII emission peak at 685 nm. While WT, *ftsh* and *R207K^b^*^6^ show an increased PSI emission at 715 nm in reducing conditions, the *stt7-1*, *xL215^b^*^6^ and *G216^b^*^6^ mutants did not.

**Figure S5.**
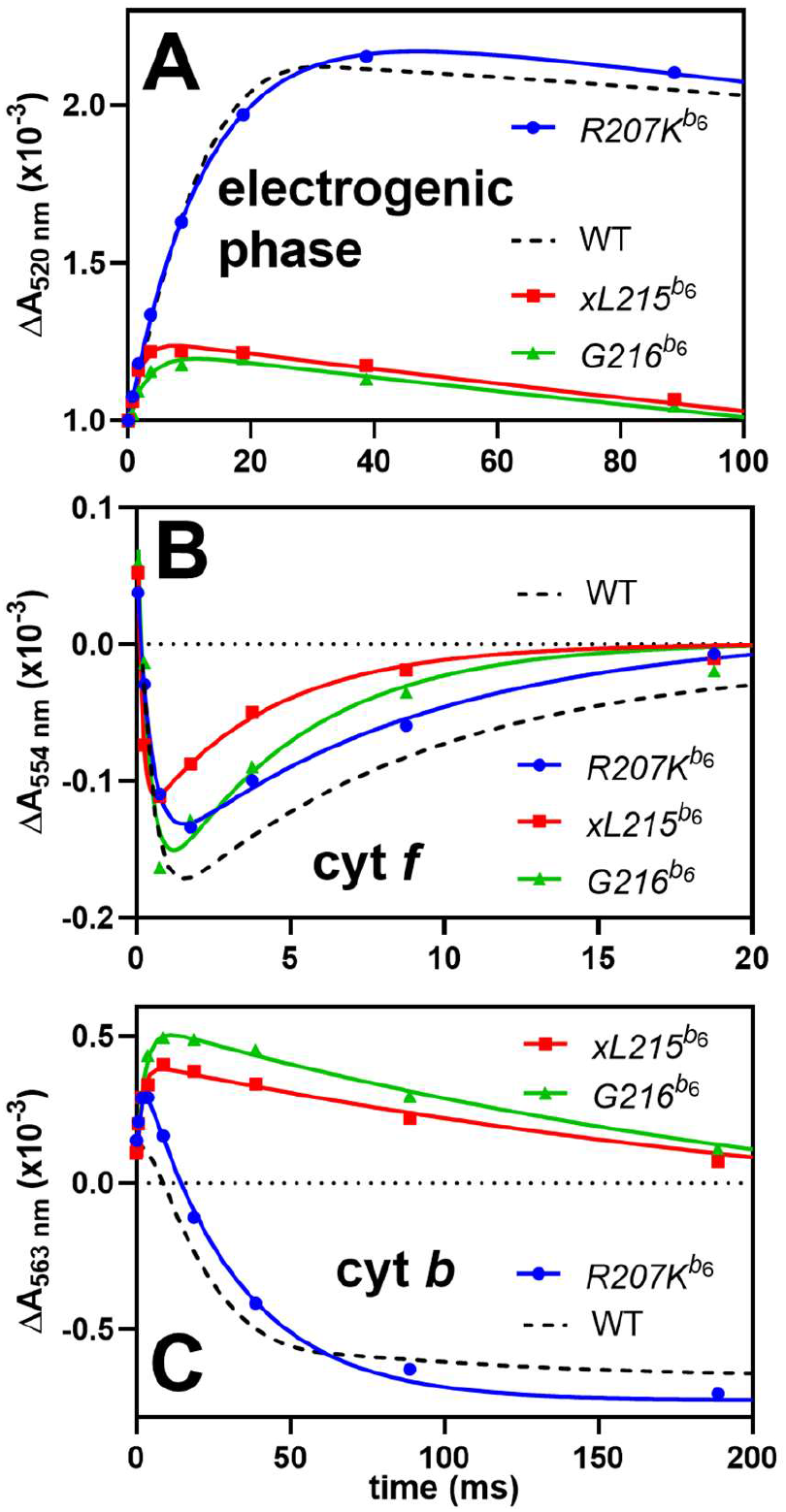
Kinetics of electron transfer in modified cyt *b*_6_*f* complexes, monitored in anaerobic intact cells treated with 10 µM DCMU and 1 mM hydroxylamine, using flash induced absorbance changes (for methods, see S7). (A) Electrogenic phase (electrochromic shift at 520 nm) showing the Q_o_H_2_ to Q_i_ electron transfer (Q-cycle), through the “low-potential chain” of cofactors *b*_L_, *b*_H_, and *c*_i_. While this phase is not affected in *R207K^b^*^6^, it is strongly reduced in *xL215^b^*^6^ and *G216^b^*^6^ mutants. (B) Cytochrome *f* oxidation by plastocyanin (fast absorbance decrease at 554 nm) and reduction (slower increase) by Q_o_ shows that the “high-potential chain” is not affected. (C) Cytochrome *b*_L_ reduction by Q_o_, (fast absorbance increase at 563 nm) and oxidation of *b*_L_ and *b*_H_ (slower decrease) leading to a net oxidation of *b*_H_ (reduced in the dark prior to the flash under anoxia) in *R207K^b^*^6^. In *xL215^b^*^6^ and *G216^b^*^6^ while cyt *b* reduction from Q_o_ is unchanged, cyt *b* oxidation by Q_i_ is drastically slowed down due to the absence of heme *c*_i_.

**Text S1. Methods used for *in vivo* time-resolved spectrophotometry** used in Figure S6. Flash-induced absorbance changes were monitored using a JTS10 spectrophotometer (BioLogic) upgraded by Daniel Béal (JBeamBio) where the light source is a pulsed light-emitting diode and the measuring wavelength is selected through various interference filters. Single turnover sub-saturating flashes are provided by a pulsed Nd:YAG laser pumping an optical parametric oscillator (Continuum lasers). For a guide of good practice using this apparatus, see the recent work of Mathiot and Alric ^9^. Treatment with 10 µM DCMU and 1 mM hydroxylamine fully inhibited Photosytem II after a preillumination. The actinic flash instantly oxidizes P700, then Plastocyanin (few microseconds), then cyt *f* (few hundreds of microseconds) / Rieske Iron-Sulfur Protein, then the quinol (QH_2_) bound in Q_o_. Therefore, the cyt *f* reduction phase (∼3 ms at 554 nm) corresponds to quinol oxidation. The second electron from the quinol is directed towards the low potential chain and transferred to heme *b*_L_ and heme *b*_H_. At room temperature, these hemes are very close in absorbance and barely distinguishable ^10^. In practice, after > 1 hour dark anaerobic adaptation, *b*_H_ becomes reduced, due to the reducing poise inside the cells. Following a PSI flash excitation, the injection of an electron in the low potential chain therefore reduces *b*_L_, giving a small transient increase at 563 nm, while both *b*_H_ and *b*_L_ are being oxidized by the quinone in Q_i_, observable as a net decrease in absorbance. After the flash, a long-lived oxidized *b* heme signal is taken as an internal control that *b*_H_ was reduced prior to the flash. Repetitive flashes increase the reduction phase and decrease the net oxidation, showing the randomization of the redox state of the *b* hemes through multiple Q-cycles ^11^.

**Text S2. Protein sequence alignment of STT/N7 proteins** from Micromonas (*Mc*) Chlamydomonas (*Cr*) or Arabidopsis (*At*). Residues found in their phosphorylated form in this work are shown in red and pink. Important protein domains for kinase activation are shown in green. The kinase domain is in bold.

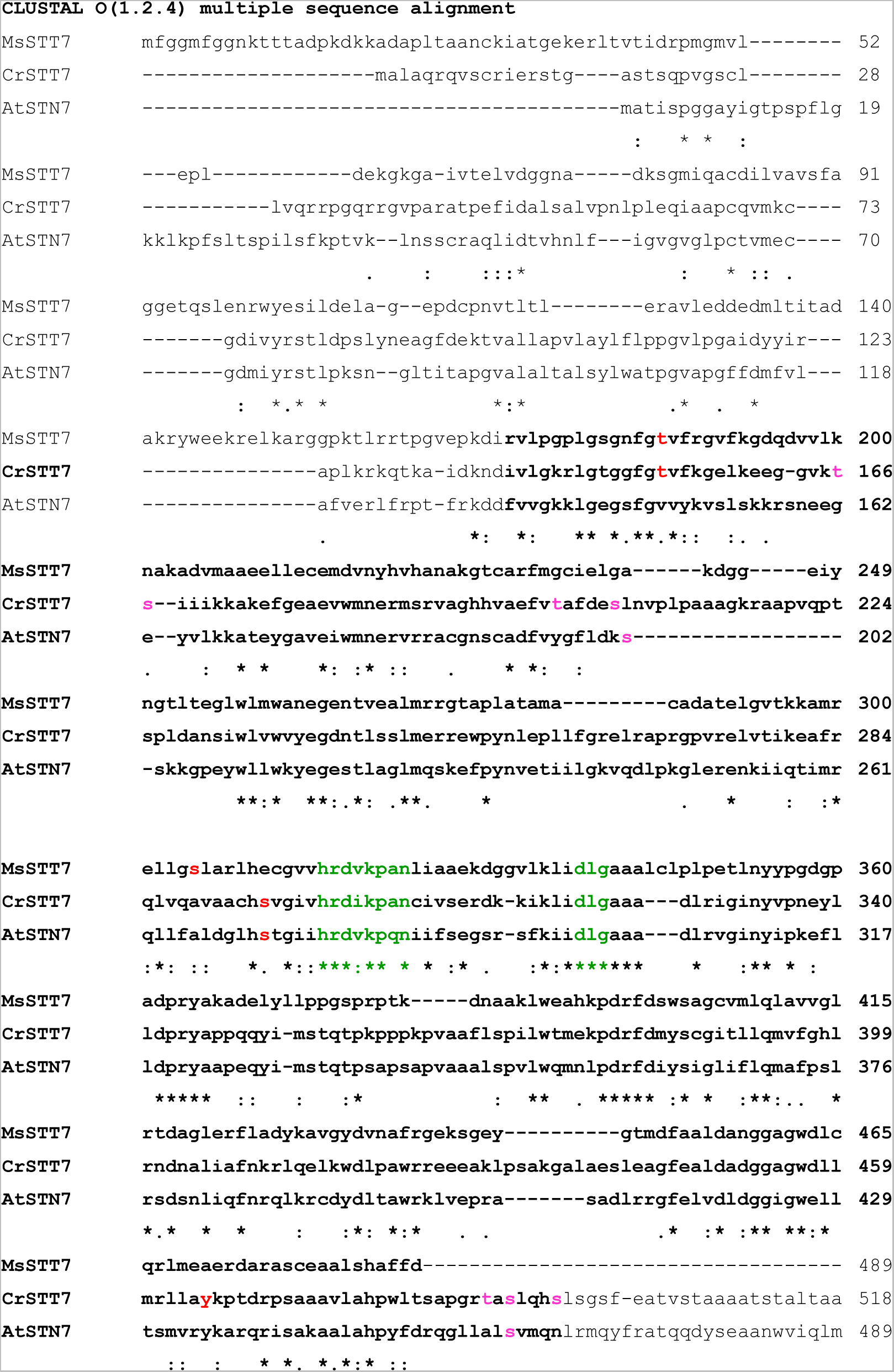
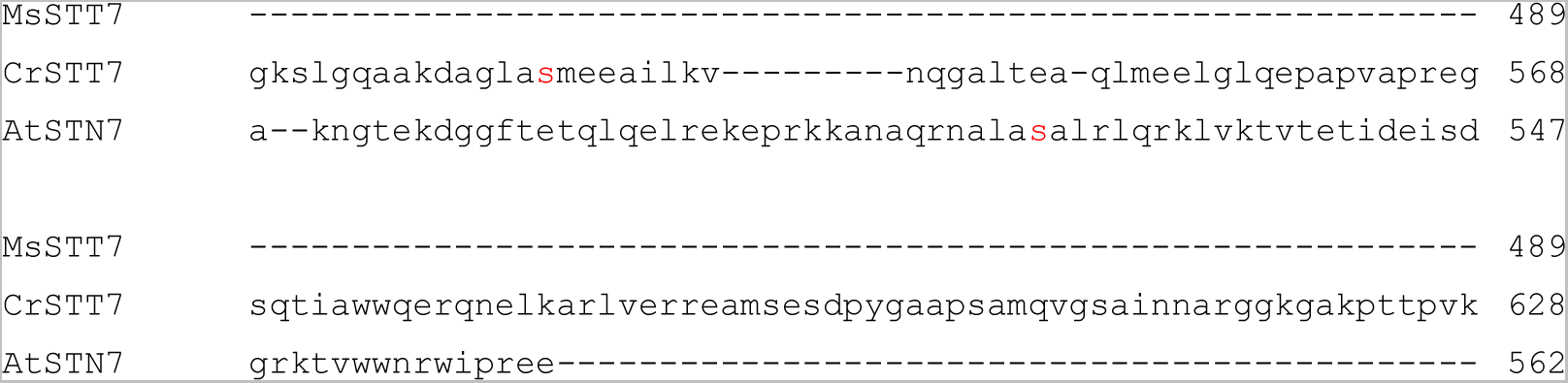

## References

1. Depège, N., Bellafiore, S. & Rochaix, J.-D. Role of Chloroplast Protein Kinase Stt7 in LHCII Phosphorylation and State Transition in Chlamydomonas. Science (2003).

2. Cariti, F. et al. Regulation of Light Harvesting in Chlamydomonas reinhardtii Two Protein Phosphatases Are Involved in State Transitions. Plant Physiol. 183, 1749–1764 (2020).

3. Lemaire, C., Girard-Bascou, J., Wollman, F.-A. & Bennoun, P. Studies on the cytochrome b6/f complex. I. Characterization of the complex subunits in Chlamydomonas reinhardtii. Biochim. Biophys. Acta BBA - Bioenerg. 851, 229–238 (1986).

4. Vener, A. V., Van Kan, P. J., Gal, A., Andersson, B. & Ohad, I. Activation/deactivation cycle of redox-controlled thylakoid protein phosphorylation. Role of plastoquinol bound to the reduced cytochrome bf complex. J. Biol. Chem. 270, 25225–25232 (1995).

5. Zito, F. et al. The Qo site of cytochrome b6f complexes controls the activation of the LHCII kinase. EMBO J. 18, 2961–2969 (1999).

6. Finazzi, G., Zito, F., Barbagallo, R. P. & Wollman, F. A. Contrasted effects of inhibitors of cytochrome b6f complex on state transitions in Chlamydomonas reinhardtii: the role of Qo site occupancy in LHCII kinase activation. J. Biol. Chem. 276, 9770–9774 (2001).

7. Lemeille, S. et al. Analysis of the Chloroplast Protein Kinase Stt7 during State Transitions. PLoS Biol. 7, e45 (2009).

8. Shapiguzov, A. et al. Activation of the Stt7/STN7 Kinase through Dynamic Interactions with the Cytochrome b6f Complex. Plant Physiol. 171, 82–92 (2016).

9. Dumas, L. et al. A stromal region of cytochrome b6f subunit IV is involved in the activation of the Stt7 kinase in Chlamydomonas. Proc. Natl. Acad. Sci. 114, 12063–12068 (2017).

10. Stroebel, D., Choquet, Y., Popot, J.-L. & Picot, D. An atypical haem in the cytochrome b(6)f complex. Nature 426, 413–418 (2003).

11. Malnoë, A., Wang, F., Girard-Bascou, J., Wollman, F.-A. & de Vitry, C. Thylakoid FtsH protease contributes to photosystem II and cytochrome b6f remodeling in Chlamydomonas reinhardtii under stress conditions. Plant Cell 26, 373–390 (2014).

12. Kuras, R. & Wollman, F. A. The assembly of cytochrome b6/f complexes: an approach using genetic transformation of the green alga Chlamydomonas reinhardtii. EMBO J. 13, 1019–1027 (1994).

13. Kuras, R. et al. Molecular genetic identification of a pathway for heme binding to cytochrome b6. J. Biol. Chem. 272, 32427–32435 (1997).

14. Bergner, S. V. et al. STATE TRANSITION7-Dependent Phosphorylation Is Modulated by Changing Environmental Conditions, and Its Absence Triggers Remodeling of Photosynthetic Protein Complexes1. Plant Physiol. 168, 615–634 (2015).

15. Hamel, P., Olive, J., Pierre, Y., Wollman, F. A. & de Vitry, C. A new subunit of cytochrome b6f complex undergoes reversible phosphorylation upon state transition. J. Biol. Chem. 275, 17072–17079 (2000).

16. Buchert, F. et al. The labile interactions of cyclic electron flow effector proteins. J. Biol. Chem. jbc.RA118.004475 (2018) doi:10.1074/jbc.RA118.004475.

17. Chen, X., Kindle, K. & Stern, D. Initiation codon mutations in the Chlamydomonas chloroplast petD gene result in temperature-sensitive photosynthetic growth. EMBO J. 12, 3627–3635 (1993).

18. Zito, F., Vinh, J., Popot, J.-L. & Finazzi, G. Chimeric Fusions of Subunit IV and PetL in the b6 f Complex of Chlamydomonas reinhardtii: STRUCTURAL IMPLICATIONS AND CONSEQUENCES ON STATE TRANSITIONS*. J. Biol. Chem. 277, 12446–12455 (2002).

19. Lemeille, S., Turkina, M. V., Vener, A. V. & Rochaix, J.-D. Stt7-dependent phosphorylation during state transitions in the green alga Chlamydomonas reinhardtii. Mol. Cell. Proteomics MCP 9, 1281–1295 (2010).

20. Willig, A., Shapiguzov, A., Goldschmidt-Clermont, M. & Rochaix, J.-D. The phosphorylation status of the chloroplast protein kinase STN7 of Arabidopsis affects its turnover. Plant Physiol. 157, 2102–2107 (2011).

21. Schütz, M. et al. Early Evolution of Cytochrome bc Complexes. J. Mol. Biol. 300, 663–675 (2000).

22. Kuras, R., Saint-Marcoux, D., Wollman, F.-A. & de Vitry, C. A specific c-type cytochrome maturation system is required for oxygenic photosynthesis. Proc. Natl. Acad. Sci. 104, 9906–9910 (2007).

23. Saint-Marcoux, D., Wollman, F.-A. & de Vitry, C. Biogenesis of cytochrome b6 in photosynthetic membranes. J. Cell Biol. 185, 1195–1207 (2009).

24. Malnoë, A., Wollman, F.-A., de Vitry, C. & Rappaport, F. Photosynthetic growth despite a broken Q-cycle. Nat. Commun. 2, 301 (2011).

25. Takahashi, H. et al. PETO Interacts with Other Effectors of Cyclic Electron Flow in Chlamydomonas. Mol. Plant 9, 558–568 (2016).

26. Proctor, M. S. et al. Cryo-EM structures of the Synechocystis sp. PCC 6803 cytochrome b6f complex with and without the regulatory PetP subunit. Biochem. J. 479, 1487–1503 (2022).

27. Rexroth, S. et al. Functional Characterization of the Small Regulatory Subunit PetP from the Cytochrome b6f Complex in Thermosynechococcus elongatus. Plant Cell 26, 3435– 3448 (2014).

28. Sarewicz, M. et al. High-resolution cryo-EM structures of plant cytochrome b6f at work. Sci. Adv. 9, eadd9688 (2023).

29. Reinhardt, R. & Leonard, T. A. A critical evaluation of protein kinase regulation by activation loop autophosphorylation. eLife 12, e88210 (2023).

30. Trotta, A., Suorsa, M., Rantala, M., Lundin, B. & Aro, E.-M. Serine and threonine residues of plant STN7 kinase are differentially phosphorylated upon changing light conditions and specifically influence the activity and stability of the kinase. *Plant J*. Cell Mol. Biol. 87, 484–494 (2016).

31. Guo, J. et al. Structure of the catalytic domain of a state transition kinase homolog from Micromonas algae. Protein Cell 4, 607–619 (2013).

32. Gatti, A., Huang, Z., Tuazon, P. T. & Traugh, J. A. Multisite autophosphorylation of p21-activated protein kinase gamma-PAK as a function of activation. J. Biol. Chem. 274, 8022– 8028 (1999).

33. Perutz, M. F. Stereochemistry of Cooperative Effects in Haemoglobin: Haem–Haem Interaction and the Problem of Allostery. Nature 228, 726–734 (1970).

34. Alric, J., Pierre, Y., Picot, D., Lavergne, J. & Rappaport, F. Spectral and redox characterization of the heme ci of the cytochrome b6f complex. Proc. Natl. Acad. Sci. 102, 15860–15865 (2005).

35. Baymann, F., Giusti, F., Picot, D. & Nitschke, W. The ci/bH moiety in the b6f complex studied by EPR: a pair of strongly interacting hemes. Proc. Natl. Acad. Sci. U. S. A. 104, 519–524 (2007).

36. Yamashita, E., Zhang, H. & Cramer, W. A. Structure of the Cytochrome b6f Complex: Quinone Analogue Inhibitors as Ligands of Heme cn. J. Mol. Biol. 370, 39–52 (2007).

37. Malone, L. A. et al. Cryo-EM structure of the spinach cytochrome b6 f complex at 3.6 Å resolution. Nature 575, 535–539 (2019).

38. Gallaher, S. D., Fitz-Gibbon, S. T., Glaesener, A. G., Pellegrini, M. & Merchant, S. S. Chlamydomonas Genome Resource for Laboratory Strains Reveals a Mosaic of Sequence Variation, Identifies True Strain Histories, and Enables Strain-Specific Studies. Plant Cell 27, 2335–2352 (2015).

39. Goldschmidt-Clermont, M. Transgenic expression of aminoglycoside adenine transferase in the chloroplast: a selectable marker of site-directed transformation of chlamydomonas. Nucleic Acids Res. 19, 4083–4089 (1991).

40. Johnson, X. et al. A new setup for in vivo fluorescence imaging of photosynthetic activity. Photosynth. Res. 102, 85–93 (2009).

41. Thomas, P. E., Ryan, D. & Levin, W. An improved staining procedure for the detection of the peroxidase activity of cytochrome P-450 on sodium dodecyl sulfate polyacrylamide gels. Anal. Biochem. 75, 168–176 (1976).

42. Cubillos-Rojas, M. et al. Tris-acetate polyacrylamide gradient gels for the simultaneous electrophoretic analysis of proteins of very high and low molecular mass. Methods Mol. Biol. Clifton NJ 869, 205–213 (2012).

43. Kelley, L. A., Mezulis, S., Yates, C. M., Wass, M. N. & Sternberg, M. J. E. The Phyre2 web portal for protein modeling, prediction and analysis. Nat. Protoc. 10, 845–858 (2015).

## References

2. Bergner, S. V. et al. STATE TRANSITION7-Dependent Phosphorylation Is Modulated by Changing Environmental Conditions, and Its Absence Triggers Remodeling of Photosynthetic Protein Complexes1. Plant Physiol. 168, 615–634 (2015).

3. Kuras, R. & Wollman, F. A. The assembly of cytochrome b6/f complexes: an approach using genetic transformation of the green alga Chlamydomonas reinhardtii. EMBO J. 13, 1019–1027 (1994).

4. Chen, X., Kindle, K. & Stern, D. Initiation codon mutations in the Chlamydomonas chloroplast petD gene result in temperature-sensitive photosynthetic growth. EMBO J. 12, 3627–3635 (1993).

5. Zito, F., Vinh, J., Popot, J.-L. & Finazzi, G. Chimeric Fusions of Subunit IV and PetL in the b6 f Complex of Chlamydomonas reinhardtii: STRUCTURAL IMPLICATIONS AND CONSEQUENCES ON STATE TRANSITIONS*. J. Biol. Chem. 277, 12446–12455 (2002).

6. Zito, F. et al. The Qo site of cytochrome b6f complexes controls the activation of the LHCII kinase. EMBO J. 18, 2961–2969 (1999).

7. Dumas, L. et al. A stromal region of cytochrome b6f subunit IV is involved in the activation of the Stt7 kinase in Chlamydomonas. Proc. Natl. Acad. Sci. 114, 12063–12068 (2017).

8. Malnoë, A., Wang, F., Girard-Bascou, J., Wollman, F.-A. & de Vitry, C. Thylakoid FtsH protease contributes to photosystem II and cytochrome b6f remodeling in Chlamydomonas reinhardtii under stress conditions. Plant Cell 26, 373–390 (2014).

9. Mathiot, C. & Alric, J. Standard units for ElectroChromic Shift measurements in plant biology. J. Exp. Bot. 72, 6467–6473 (2021).

10. Joliot, P. & Joliot, A. The low-potential electron-transfer chain in the cytochrome bf complex. Biochim. Biophys. Acta BBA - Bioenerg. 933, 319–333 (1988).

11. Joliot, P. & Joliot, A. Electrogenic events associated with electron and proton transfers within the cytochrome b(6)/f complex. Biochim. Biophys. Acta 1503, 369–376 (2001).

